# Virucidal and antiviral activity of astodrimer sodium against SARS-CoV-2 *in vitro*

**DOI:** 10.1101/2020.08.20.260190

**Authors:** Jeremy R.A. Paull, Graham P. Heery, Michael D. Bobardt, Alex Castellarnau, Carolyn A. Luscombe, Jacinth K. Fairley, Philippe A. Gallay

**Affiliations:** Starpharma Pty Ltd, 4-6 Southampton Crescent, Abbotsford, Victoria 3067, Australia; Department of Immunology and Microbiology, The Scripps Research Institute, La Jolla, CA 92307, USA

**Keywords:** Astodrimer, COVID-19, dendrimer, antiviral, SARS-CoV-2, SPL7013

## Abstract

An effective response to the ongoing coronavirus disease (COVID-19) pandemic caused by severe acute respiratory syndrome coronavirus 2 (SARS-CoV-2) will involve a range of complementary preventive modalities. The current studies were conducted to evaluate the *in vitro* SARS-CoV-2 antiviral and virucidal activity of astodrimer sodium, a dendrimer with broad spectrum antimicrobial activity, including against enveloped viruses in *in vitro* and *in vivo* models, that is marketed for antiviral and antibacterial applications. We report that astodrimer sodium inhibits replication of SARS-CoV-2 in Vero E6 and Calu-3 cells, with 50% effective concentrations (EC_50_) for i) reducing virus-induced cytopathic effect of 0.002 to 0.012 mg/mL in Vero E6 cells, and ii) infectious virus release by plaque assay of 0.019 to 0.031 mg/mL in Vero E6 cells and 0.031 to 0.045 mg/mL in Calu-3 cells. The selectivity index (SI) in these assays was as high as 2197. Astodrimer sodium was also virucidal, reducing SARS-CoV-2 infectivity by >99.9% (>3 log_10_) within 1 minute of exposure, and up to >99.999% (>5 log_10_) shown at astodrimer sodium concentrations of 10 to 30 mg/mL in Vero E6 and Calu-3 cell lines. Astodrimer sodium also inhibited infection in a primary human airway epithelial cell line. The data were similar for all investigations and were consistent with the potent antiviral and virucidal activity of astodrimer sodium being due to inhibition of virus-host cell interactions, as previously demonstrated for other viruses. Further studies will confirm if astodrimer sodium binds to SARS-CoV-2 spike protein and physically blocks initial attachment of the virus to the host cell. Given the *in vitro* effectiveness and significantly high SI, astodrimer sodium warrants further investigation for potential as a nasally administered or inhaled antiviral agent for SARS-CoV-2 prevention and treatment applications.

## 1. Introduction

The ongoing pandemic coronavirus disease 2019 (COVID-19), caused by severe acute respiratory syndrome coronavirus-2 (SARS-CoV-2) infection, has resulted in unprecedented efforts to rapidly develop strategies to contain infection rates for the protection of vulnerable populations. An effective public health response to the current pandemic will involve currently available vaccines being complemented by supplementary preventive modalities.

SARS-CoV-2 receptors and coreceptors have been shown to be highly expressed in nasal epithelial cells (Sungnak et al., 2020). This finding is consistent with the virus infectivity or replication pattern along the respiratory tract, which peaks proximally (nasal cavity) and is relatively minimal in the distal alveolar regions (Hou et al., 2020). These findings suggest that nasal carriage of the virus is a key feature of transmission, and that nasally administered therapeutic modalities could be potentially effective in helping to prevent spread of infection.

Astodrimer sodium (SPL7013) is a generation-four lysine dendrimer with a polyanionic surface charge (McCarthy et al., 2005) that is active against several enveloped and non-enveloped viruses including human immunodeficiency virus-1 (HIV-1) (Lackman-Smith et al., 2008, Tyssen et al., 2010), herpes simplex virus (HSV)−1 and −2 (Gong et al., 2005), H1N1 and H3N2 influenza virus, human respiratory syncytial virus (HRSV), human papillomavirus (HPV), adenovirus and Zika virus (unpublished data). Astodrimer sodium also has antibacterial properties. Both size and surface charge contribute to the function of the compound (Tyssen et al., 2010), and when administered topically, astodrimer sodium is not absorbed systemically (Chen et al., 2009; O’Loughlin et al., 2010; McGowan et al., 2011).

Vaginally administered astodrimer sodium protected macaques from infection with chimeric simian-HIV-1 (SHIV)_89.6P_ (Jiang et al., 2005), and mice and guinea pigs from HSV-2 infection (Bernstein et al., 2003) in vaginal infection challenge models. Astodrimer 1% Gel (10 mg/mL astodrimer sodium) administered vaginally has been shown to be safe and effective in phase 2 and large phase 3 trials for treatment and prevention of bacterial vaginosis (BV) (Chavoustie et al., 2020; Waldbaum et al., 2020; Schwebke et al., 2021) and is marketed in Europe, Australia, New Zealand and several countries in Asia.

The current studies were conducted to assess the antiviral and virucidal activity of astodrimer sodium against SARS-CoV-2 *in vitro*, to determine its potential as a reformulated, nasally administered or inhaled antiviral agent to help prevent spread of SARS-CoV-2 infection.

## 2. Materials and methods

### 2.1 Virus, cell culture, astodrimer sodium and controls

SARS-CoV-2 hCoV-19/Australia/VIC01/2020 was a gift from Melbourne’s Peter Doherty Institute for Infection and Immunity (Melbourne, Australia). Virus stock was generated at 360Biolabs (Melbourne, Australia) by two passages in Vero cells in virus growth media, which comprised Minimal Essential Medium (MEM) without L-glutamine supplemented with 1% (w/v) L-glutamine, 1.0 μg/mL of L-(tosylamido-2-phenyl) ethyl chloromethyl ketone (TPCK)-treated trypsin (Worthington Biochemical, NJ, USA), 0.2% bovine serum albumin (BSA) and 1% insulin-transferrin-selenium (ITS).

SARS-CoV-2 2019-nCoV/USA-WA1/2020 strain was isolated from an oropharyngeal swab from a patient with a respiratory illness who developed clinical disease (COVID-19) in January 2020 in Washington, US, and sourced from BEI Resources (NR-52281). Virus was derived from African green monkey kidney Vero E6 cells or lung homogenates from human angiotensin converting enzyme 2 (hACE2) transgenic mice.

Vero E6 and human Calu-3 cell lines were cultured in MEM without L-glutamine supplemented with 10% (v/v) heat-inactivated fetal bovine serum (FBS) and 1% (w/v) L-glutamine. Vero E6 and Calu-3 cells were passaged for a maximum of 10 passages for antiviral and virucidal studies. Hank’s balanced salt solution (HBSS) with 2% FBS was used for infection. The 2019-nCoV/USA-WA1/2020 strain antiviral assays were performed with a multiplicity of infection (MOI) of 0.1.

The virus inoculums for virucidal assays were 10^4^, 10^5^, and 10^6^ pfu/mL. After defined incubation periods, the solution was pelleted through a 20% sucrose cushion (Beckman SW40 Ti rotor) and resuspended in 1.5 mL MEM, which was then added to 2.5×10^4^ cells/well.

Primary human bronchial epithelial cells (HBEpC) (Sigma-Aldrich, MO, USA) were grown and maintained in HBEpC/HTEpC growth medium (Cell Applications, CA, USA). These primary cells express the ACE2 receptor and are permissive to SARS-CoV-2 infection. These cells were used to determine the antiviral effect of astodrimer sodium against SARS-CoV-2 in a primary human airway epithelial cell line. Cells were infected with SARS-CoV-2 2019-nCoV/USA-WA1/2020 at 10^3^ pfu/mL with 1 mL added to 2.5×10^4^ cells/well. The positive control was addition of 10 μg/mL of SARS-CoV-2 spike protein antibody (pAb, T01KHuRb) (ThermoFisher, MA, USA) at the time of infection.

Astodrimer sodium was prepared as 100 mg/mL in water and stored at 4°C. Astodrimer sodium has a molecular weight of 16581.57 g/mol and the structure is described and illustrated in Tyssen et al., 2010. The purity of the compound used in these studies was assessed by ultra-high-performance liquid chromatography (UPLC) to be 98.79%.

Remdesivir (MedChemExpress, NJ, USA) was used as a positive control in the virus-induced cytopathic effect (CPE) inhibition and plaque assays.

Iota-carrageenan (Sigma-Aldrich, MO, USA) was used in the primary epithelial cell nucleocapsid and plaque assays to compare the antiviral activity of this substance with astodrimer sodium. Concentrations used are those reported to show activity against SARS-CoV-2 (Bansal et al., 2020).

### 2.2 Virus-induced cytopathic effect inhibition assay

Vero E6 (ATCC-CRL1586) cell stocks were generated in cell growth medium, which comprised MEM without L-glutamine supplemented with 10% (v/v) heat-inactivated FBS and 1% (w/v) L-glutamine. Vero E6 cell monolayers were seeded in 96-well plates at 2×10^4^ cells/well in 100 μL growth medium (MEM supplemented with 1% (w/v) L-glutamine, 2% FBS) and incubated overnight at 37°C in 5% CO_2_. SARS-CoV-2 infection was established by using an MOI of 0.05 to infect cell monolayers.

Astodrimer sodium or remdesivir were serially diluted 1:3, 9 times and each compound concentration was assessed for both antiviral efficacy and cytotoxicity in triplicate. Astodrimer sodium was added to Vero E6 cells 1 hour prior to infection or 1 hour post-infection with SARS-CoV-2. Cell cultures were incubated at 37°C in 5% CO_2_ for 4 days prior to assessment of CPE. The virus growth media was MEM supplemented with 1% (w/v) L-glutamine, 2% FBS, and 4 μg/mL TPCK-treated trypsin. On Day 4, viral-induced CPE and cytotoxicity of the compound were determined by measuring the viable cells using the methylthiazolyldiphenyl-tetrazolium bromide (MTT) assay (MP Biomedicals, NSW, Australia). Absorbance was measured at 540-650 nm on a plate reader.

### 2.3 Antiviral plaque assay evaluation and nucleocapsid ELISA

For the antiviral evaluation, astodrimer sodium was added to cells 1 hour prior to, at the time of, and 1 hour after exposing the cells to virus. For both the antiviral and virucidal assays, at 6 hours after infection, cells were washed to remove astodrimer sodium and/or any virus remaining in the supernatant, in such way that a

Following initial infection, cell cultures were incubated and supernatants recovered after 16 hours or 4 days. The amount of virus in the supernatants was determined by plaque assay (plaque forming unit [pfu]) and by nucleocapsid enzyme-linked immunosorbent assay (ELISA). The plaque assay used was as described in van den Worm et al., 2012, utilizing 2% sodium carboxymethyl cellulose overlay, fixation of cells by 4% paraformaldehyde and staining with 0.1% crystal violet. The nucleocapsid ELISA assay was as described by Bioss Antibodies, USA (BSKV0001).

The assessment of astodrimer sodium cytotoxicity occurred on Day 4 by measuring lactate dehydrogenase (LDH) activity in the cytoplasm using an LDH detection kit (Cayman Chemical), with 0.5% saponin used as the positive cytotoxic control.

### 2.4 Virucidal assay

For the virucidal evaluation, concentrations of astodrimer sodium (0.0046 to 30 mg/mL) were incubated with SARS-CoV-2 2019-nCoV/USA-WA1/2020 for times ranging from 5 seconds to 2 hours. To neutralize the effect of astodrimer sodium, unbound compound was separated from the astodrimer:virus mixture by pelleting the preincubated mixture through a 20% sucrose cushion (Beckman SW40 Ti rotor). The astodrimer sodium-containing supernatant was removed (i.e., neutralising the effect of SPL7013) and then the pelleted virus was gently resuspended and added to Vero E6 or Calu-3 cell cultures. Virus infection, cell culture and cytotoxicity assessment was as described for the plaque assay described in Section 2.3.

All antiviral and virucidal assays were performed in triplicate, except where indicated in the results.

### 2.5 Determination of 50% effective concentration (EC_50_) and cytotoxicity (CC_50_)

The concentration of compound that gives a 50% reduction in viral-induced CPE, infectious virus (pfu/mL), or secreted viral nucleocapsid (EC_50_) was calculated using the formula of Pauwels et al., 1998.

The concentration of compound that resulted in a 50% reduction in cell viability (CC_50_) after 4 days of culture was also calculated by the formula of Pauwels et al., 1998.

### 2.6 Primary Epithelial Cell Assay

Astodrimer sodium (0, 1.1, 3.3 and 10 mg/mL) or iota-carrageenan (0, 6, 60 and 600 μg/mL) were added to HBEpC cells 1 hour prior to infection with SARS-CoV-2. Cells were cultured for 4 days post-infection and the cell supernatant was analysed for the amount of secreted SARS-CoV-2 nucleocapsid by ELISA, and infectious virus was quantitated by plaque assay, as described in Section 2.3.

## 3. Results

### 3.1 Virus-induced cytopathic effect inhibition

In two independent virus-induced CPE inhibition assays, astodrimer sodium inhibited SARS-CoV-2 (hCoV-19/Australia/VIC01/2020) replication in Vero E6 cells in a dose dependent manner (Table 1). Astodrimer sodium inhibited viral replication when added either 1 hour prior to infection, or 1 hour post-infection with SARS-CoV-2.

**Table 1:**
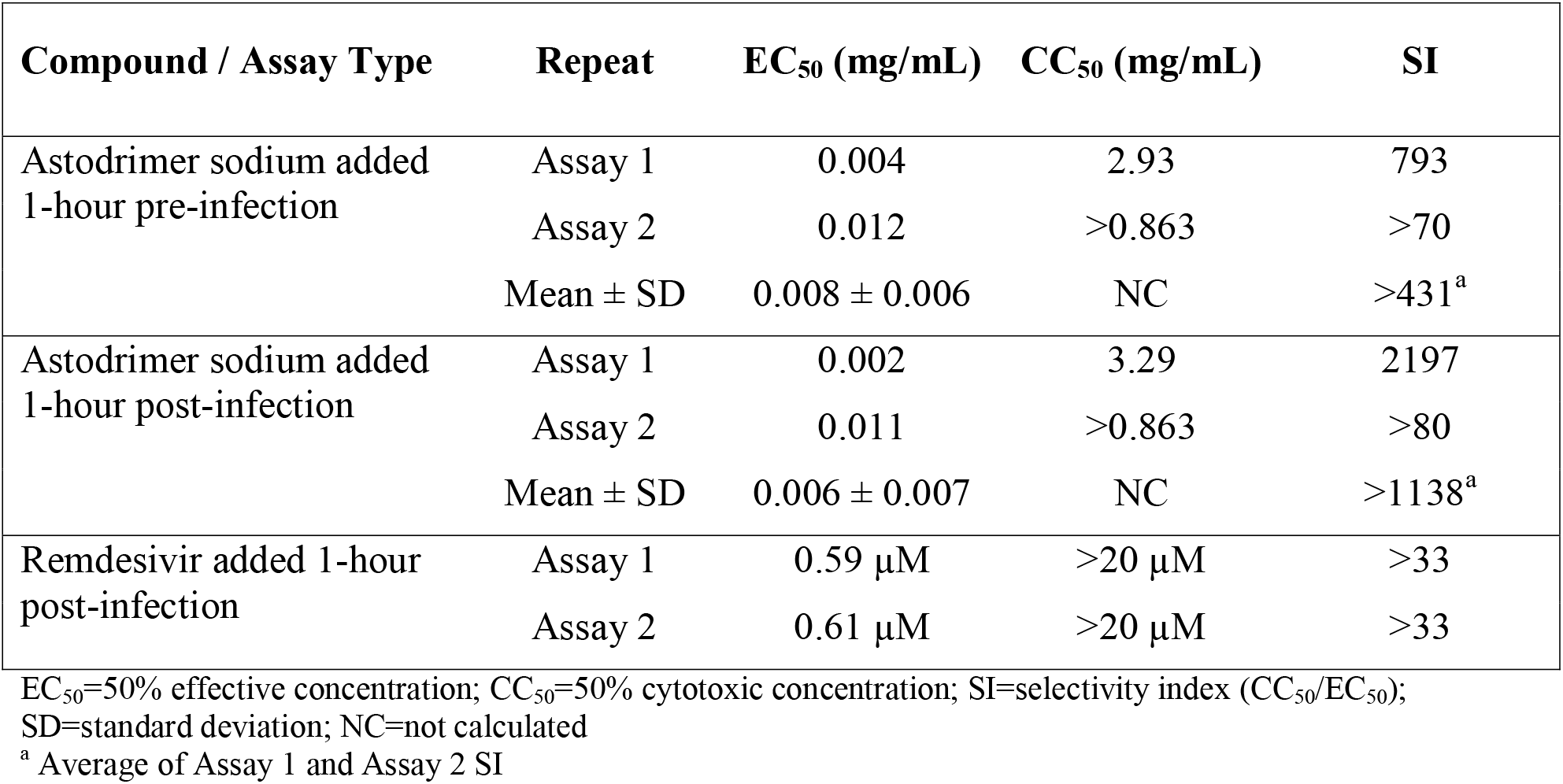
Antiviral efficacy, measured by a reduction in CPE in virus-infected cells at Day 4 post-infection, and selectivity of astodrimer sodium against SARS-CoV-2 (hCoV-19/Australia/VIC01/2020) infection of Vero E6 cells.

| Compound / Assay Type | Repeat | EC<sub>50</sub> (mg/mL) | CC<sub>50</sub> (mg/mL) | SI |
| --- | --- | --- | --- | --- |
| Astodrimmer sodium added<br>1-hour pre-infection | Assay 1 | 0.004 | 2.93 | 793 |
|  | Assay 2 | 0.012 | >0.863 | >70 |
|  | Mean ± SD | 0.008 ± 0.006 | NC | >431<sup>a</sup> |
| Astodrimmer sodium added<br>1-hour post-infection | Assay 1 | 0.002 | 3.29 | 2197 |
|  | Assay 2 | 0.011 | >0.863 | >80 |
|  | Mean ± SD | 0.006 ± 0.007 | NC | >1138<sup>a</sup> |
| Remdesivir added 1-hour<br>post-infection | Assay 1 | 0.59 µM | >20 µM | >33 |
|  | Assay 2 | 0.61 µM | >20 µM | >33 |

EC<sub>50</sub>=50% effective concentration; CC<sub>50</sub>=50% cytotoxic concentration; SI=selectivity index (CC<sub>50</sub>/EC<sub>50</sub>);
SD=standard deviation; NC=not calculated
<sup>a</sup> Average of Assay 1 and Assay 2 SI

Astodrimer sodium was initially tested in the range of 0.0013 to 8.63 mg/mL. In the repeat set of assays, astodrimer sodium was tested in the range of 0.0001 to 0.86 mg/mL to help further characterize the lower end of the dose response curve. The effective and cytotoxic concentrations, and selectivity indices from the assays are shown individually and as means in Table 1 for CPE determined readouts.

The selectivity index (SI) for astodrimer sodium against SARS-CoV-2 in the CPE studies ranged from 793 to 2197 for the initial assays where compound was added 1 hour prior to infection and 1 hour after infection, respectively, and was >70 to >80 in the repeat assays, in which cytotoxicity was not observed up to the highest concentration tested (0.86 mg/mL).

The positive control, remdesivir, was also active in the CPE inhibition assay, with a SI of >33.

### 3.2 Antiviral efficacy

To determine the ability of astodrimer sodium to inhibit globally diverse SARS-CoV-2 strains, the compound was evaluated against the 2019-nCoV/USA-WA1/2020 virus in Vero E6 cells and human Calu-3 cells. Antiviral readouts were based on virological endpoints of infectious virus or viral nucleocapsid released into the supernatant post-infection. As shown in Table 2 and Figures 1 and 2, astodrimer inhibited the 2019-nCoV/USA-WA1/2020 strain with an EC_50_ 0.019 to 0.031 mg/mL and 0.031 to 0.045 mg/mL for infectious virus release as determined by plaque assay in Vero E6 cell for Calu-3 cells, respectively. These data are consistent with the inhibition by astodrimer of the replication of the Australian SARS-CoV-2 isolate *in vitro*. The dose response data for the nucleocapsid released into the supernatant by ELISA were similar to the infectious virus release data in each cell line (data not shown). The positive control, remdesivir, was also active in the plaque assay.

**Table 2:**
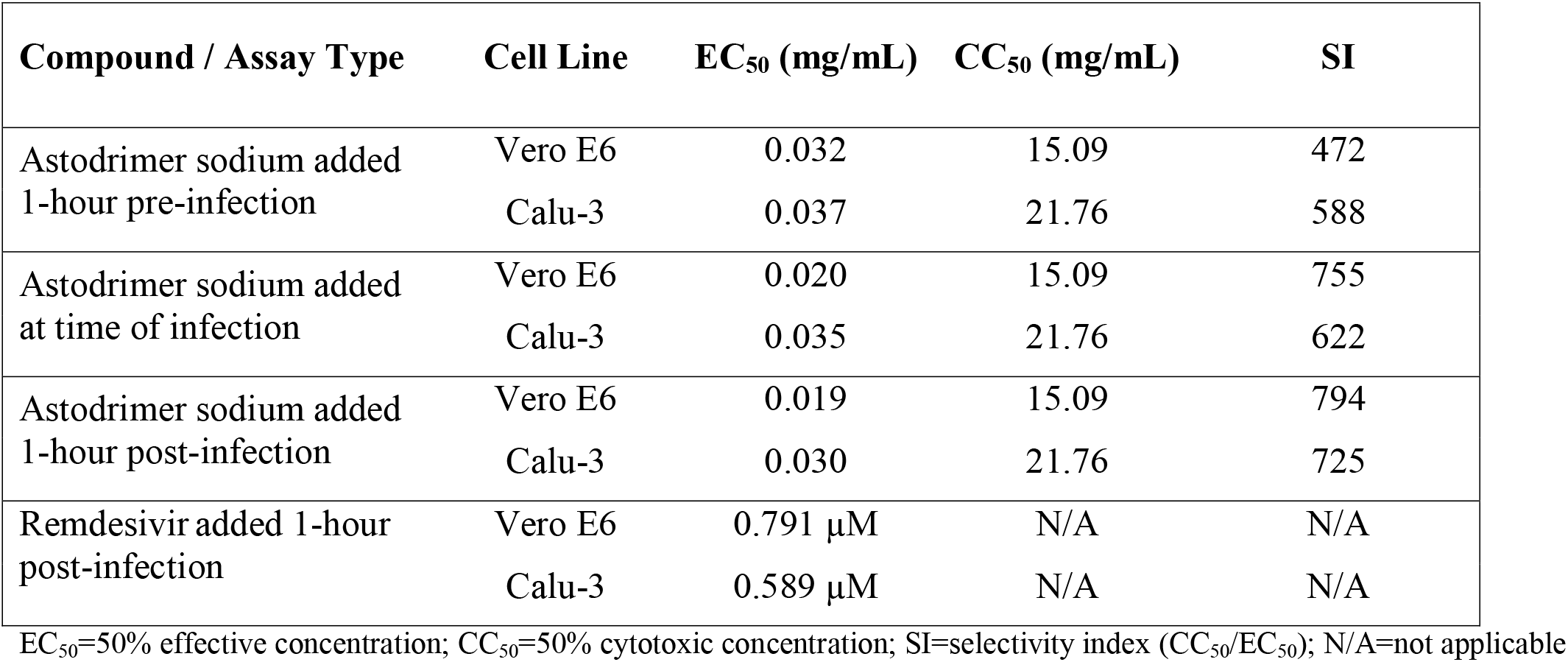
Antiviral efficacy, measured by a reduction in mean infectious virus (Log_10_ pfu/mL), and selectivity of astodrimer sodium against SARS-CoV-2 (2019-nCoV/USA-WA1/2020) on Day 4 post-infection.

**Figure 1.**
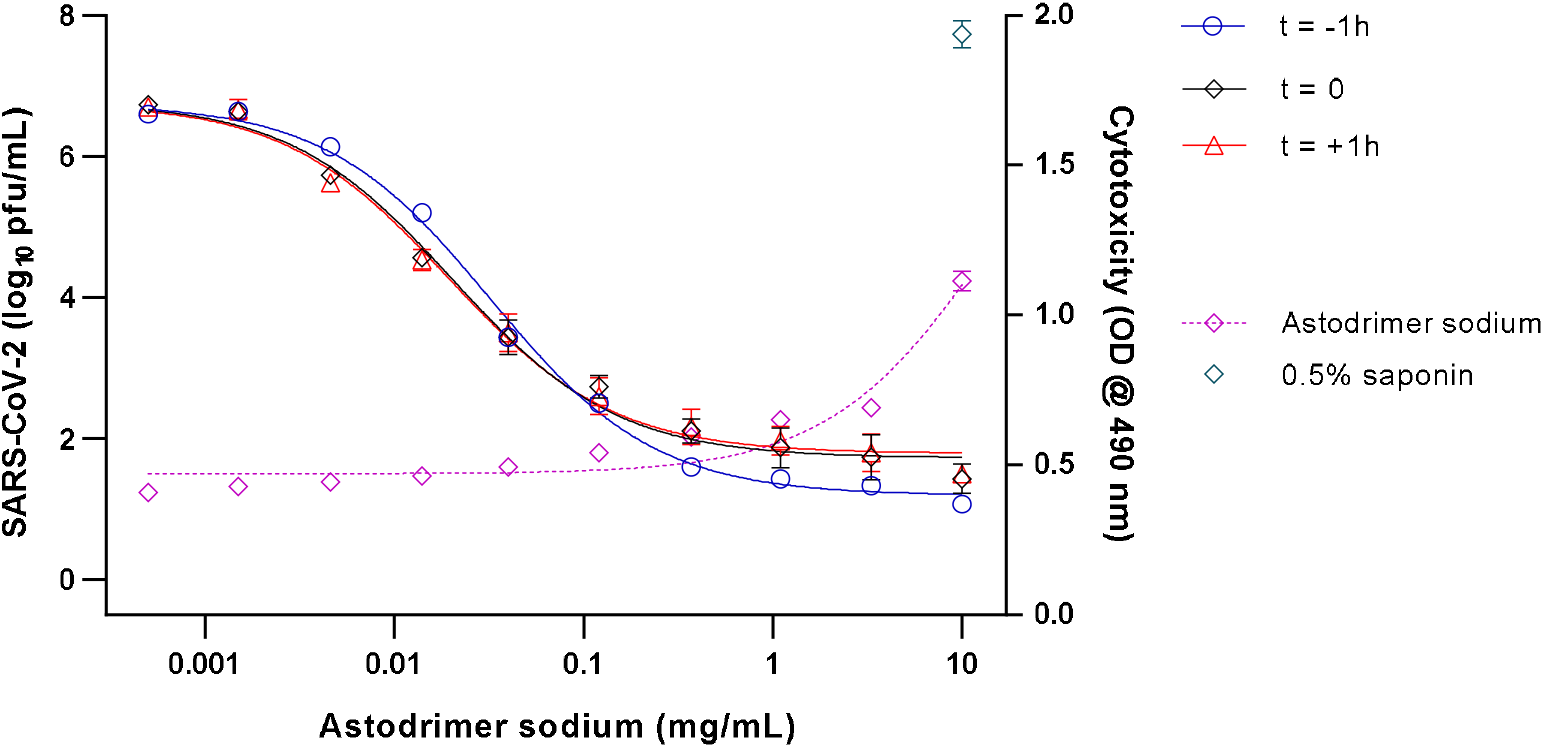
Dose-response and cytotoxicity analysis of SARS-CoV-2 (2019-nCoV/USA-WA1/2020) antiviral activity of astodrimer sodium in Vero E6 cells as measured by infectious virus release (Log_10_ pfu/mL) on Day 4 post-infection. Astodrimer sodium (0.0005 to 10 mg/mL) was added to cell cultures 1 hour prior to (t = −1h), at the time of (t = 0), and 1 hour post-infection (t = +1h). Cytotoxicity was assessed by LDH detection (OD @ 490 nm), with 0.5% saponin used as the positive cytotoxic control. Points and error bars represent mean ± SD of triplicate readings.

**Figure 2.**
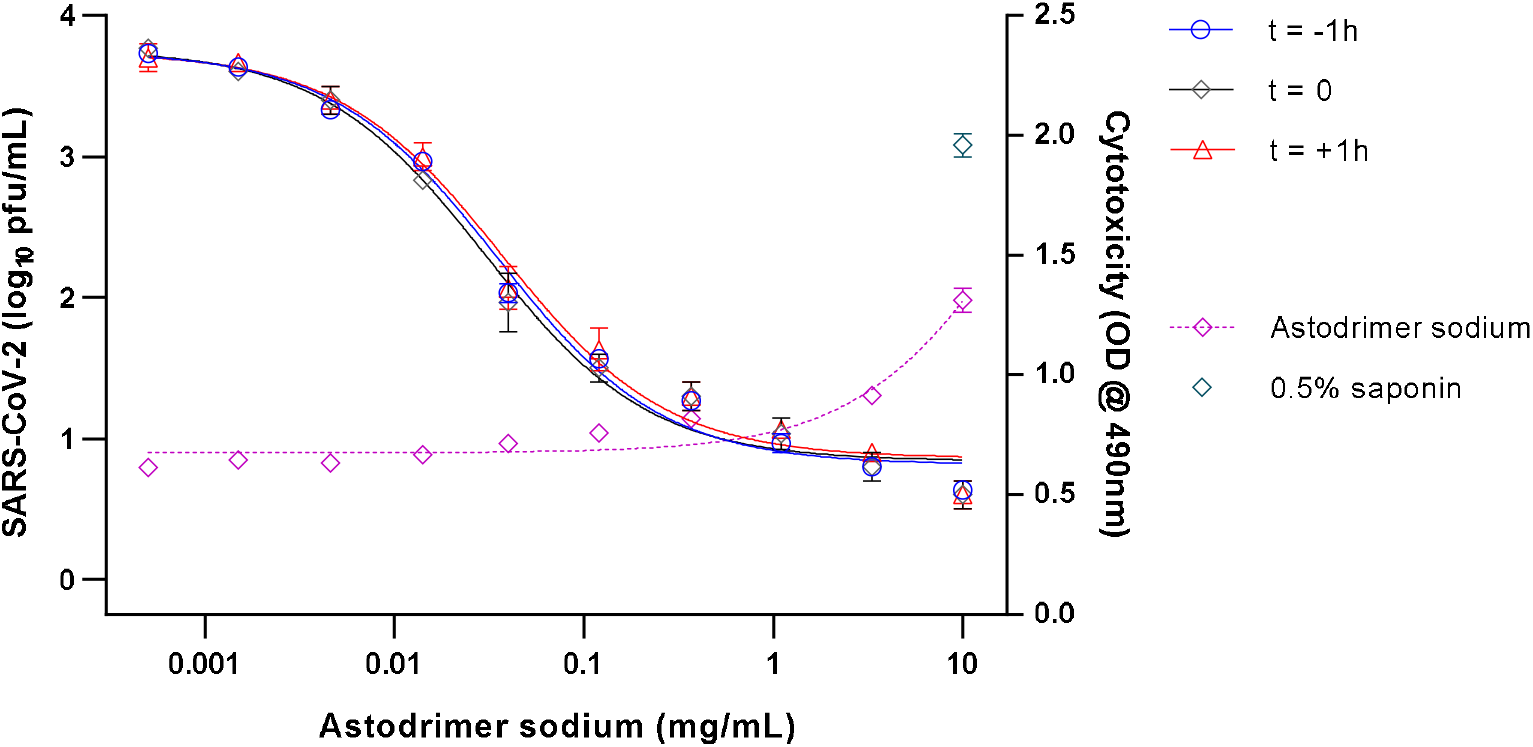
Dose-response and cytotoxicity analysis of SARS-CoV-2 (2019-nCoV/USA-WA1/2020) antiviral activity of astodrimer sodium in Calu-3 cells as measured by infectious virus release (Log_10_ pfu/mL) on Day 4 post-infection. Astodrimer sodium (0.0005 to 10 mg/mL) was added to cell cultures 1 hour prior to (t = −1h), at the time of (t Ƚ 0), and 1 hour post-infection (t = +1h). Cytotoxicity was assessed by LDH detection (OD @ 490 nm), with 0.5% saponin used as the positive cytotoxic control. Points and error bars represent mean ± SD of triplicate readings.

### 3.3 Virucidal efficacy

Virucidal assays investigated if astodrimer sodium could reduce viral infectivity by irreversibly inactivating SARS-CoV-2 prior to infection of Vero E6 cells and human airway Calu-3 cells. Following incubation of virus with astodrimer for up to 2 hours and neutralization of astodrimer, the astodrimer-exposed virus was added to cell cultures. After either 16 hours or 96 hours (Day 4), the cell culture supernatant was collected for assessment of progeny viral infectivity as determined by the amount of secreted infectious virus and nucleocapsid. The SARS-CoV-2 replication lifecycle is completed in approximately 8 hours (Ogando et al., 2020) and in these studies, we sampled at 16 hours (2 lifecycles) or Day 4 (12 lifecycles) post-infection.

Enabling a possible 12 rounds of infection, the Day 4 (96 hour) sampling time point identified that exposure of 10^6^ pfu/mL SARS-CoV-2 to astodrimer sodium for 1 to 2 hours resulted in a dose-dependent reduction in viral infectivity, with 10 to 30 mg/mL astodrimer sodium achieving up to >99.999% (>5 log_10_) reduced infectivity in Vero E6 cells and >99.9% (>3 log_10_) reduced infectivity in Calu-3 cells (data not shown) compared to untreated virus. SARS-CoV-2 infectivity was also reduced by up to >99.999% in Vero E6 cells when the incubation time of astodrimer (10 to 30 mg/mL) with 10^6^ pfu/mL virus was reduced to 15 to 30 minutes (data not shown).

Incubation of astodrimer sodium with viral inoculums of 10^4^, 10^5^ and 10^6^ pfu/mL for as little as 5 seconds resulted in evidence of reduced infectivity, with 10 to 15 minutes exposure being sufficient to achieve >99.9% reduction in virus infectivity, and greater reduction achieved with lower viral inoculum (>99.999%, 10^4^ pfu/mL viral inoculum, 10 to 30 mg/mL astodrimer sodium, and 10 to 15 min incubation time) (Table 3, Figure 3).

**Table 3:** Virucidal efficacy of 10 mg/mL astodrimer sodium against SARS-CoV-2 (2019-nCoV/USA-WA1/2020), measured by a reduction in mean infectious virus (Log_10_ pfu/mL), at 96 hours post-infection.

| Viral Load (pfu/mL) | Virus:Astodrimmer Incubation Time | Reduction vs. Virus Control ($\text{Log}_{10} \pm \text{SD}$) | Reduction vs. Virus Control (%) |
| --- | --- | --- | --- |
| $10^6$ | 5 sec | $0.10 \pm 0.20$ | 20.567 |
| | 10 sec | $0.03 \pm 0.06$ | 7.388 |
| | 30 sec | $0.10 \pm 0.10$ | 20.567 |
| | 1 min | $0.33 \pm 0.12$ | 53.584 |
| | 10 min | $2.20 \pm 0.10$ | 99.369 |
| | 15 min | $3.67 \pm 0.23$ | 99.979 |
| $10^5$ | 5 sec | $0.33 \pm 0.21$ | 53.584 |
| | 10 sec | $0.23 \pm 0.06$ | 41.566 |
| | 30 sec | $0.30 \pm 0.17$ | 49.881 |
| | 1 min | $0.47 \pm 0.21$ | 65.855 |
| | 10 min | $3.70 \pm 0.26$ | 99.980 |
| | 15 min | $4.60 \pm 0.10$ | 99.998 |
| $10^4$ | 5 sec | $-0.13 \pm 0.21$ | -35.936 |
| | 10 sec | $0.07 \pm 0.29$ | 14.230 |
| | 30 sec | $0.10 \pm 0.10$ | 20.567 |
| | 1 min | $0.10 \pm 0.00$ | 20.567 |
| | 10 min | $5.07 \pm 0.25$ | >99.999 |
| | 15 min | $5.83 \pm 0.12$ | >99.999 |

Shading indicates data points where virucidal efficacy is >99.9% (3 $\text{log}_{10}$ reduction) vs. virus control; virus control=untreated virus, 0 mg/mL astodrimmer sodium; SD=standard deviation

**Figure 3.**
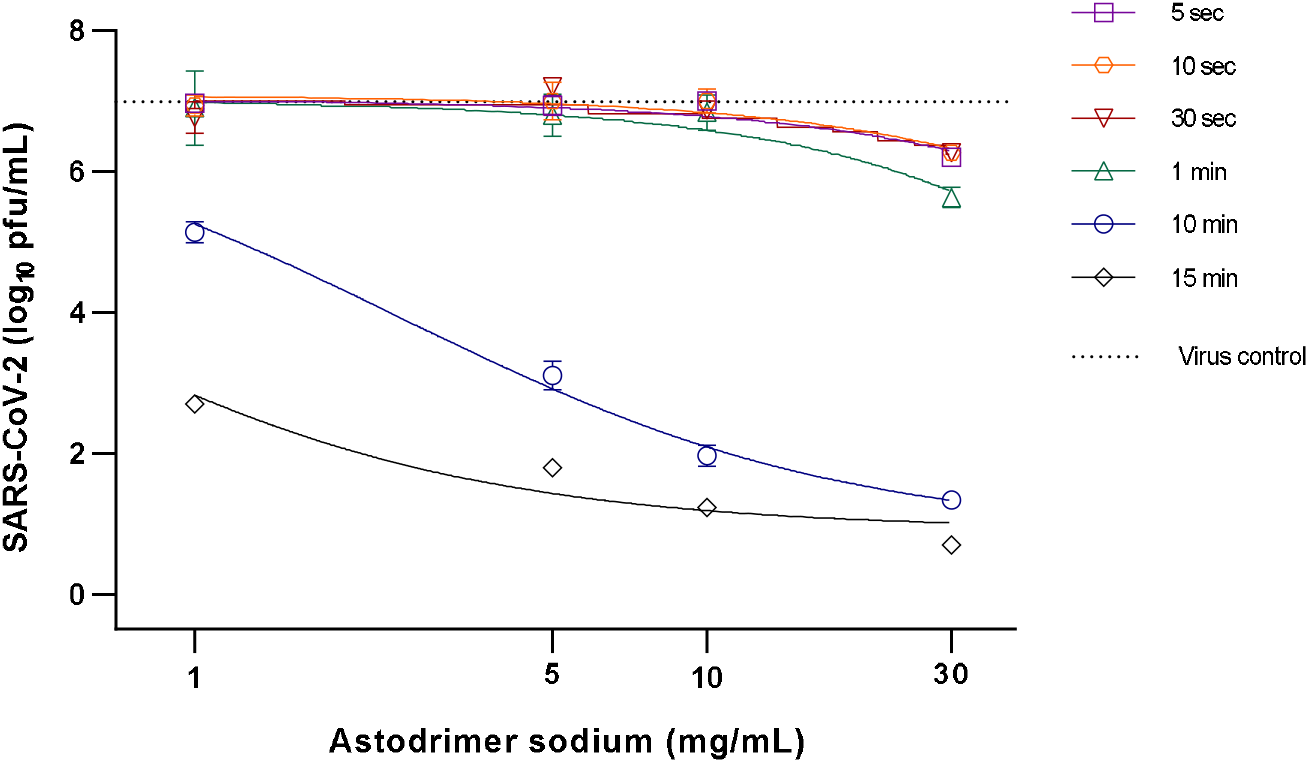
Virucidal efficacy of astodrimer sodium against SARS-CoV-2 (2019-nCoV/USA-WA1/2020) measured by a reduction in mean infectious virus (Log_10_ pfu/mL), at 96 hours post-infection in Vero E6 cells. Astodrimer sodium (1 to 30 mg/mL) was incubated with SARS-CoV-2 (2019-nCoV/USA-WA1/2020) for 5 sec up to 15 min. Treated virus was added to Vero E6 cells and the amount of infectious virus in the supernatant was determined by plaque assay 96 hours post-infection. Graph shows dose-response of astodrimer sodium virucidal activity using 10^4^ pfu/mL virus inoculum. Points and error bars represent mean ± SD of triplicate readings. Dotted line indicates level of mean infectious virus when untreated virus was added to Vero E6 cells (virus control).

When assessed 16 hours post-infection of cells with astodrimer-exposed virus, it was found that ≥10 mg/mL astodrimer sodium inactivated >99.9% SARS-CoV-2 (10^4^ pfu/mL) within as little as 1 minute of exposure (Table 4, Figure 4).

**Table 4:** Virucidal efficacy of 10 mg/mL astodrimer sodium against SARS-CoV-2 (2019-nCoV/USA-WA1/2020), measured by a reduction in mean infectious virus (Log_10_ pfu/mL), at 16 hours post-infection.

| Viral Load (pfu/mL) | Virus:Astodrimmer Incubation Time | Reduction vs. Virus Control ($\text{Log}_{10} \pm \text{SD}$) | Reduction vs. Virus Control (%) |
| --- | --- | --- | --- |
| $10^5$ | 30 sec | $0.00 \pm 0.36$ | 0.000 |
| | 1 min | $2.63 \pm 0.15$ | 99.767 |
| | 5 min | $4.63 \pm 0.31$ | 99.998 |
| | 15 min | $4.60 \pm 0.10$ | 99.998 |
| $10^4$ | 30 sec | $0.20 \pm 0.20$ | 36.904 |
| | 1 min | $3.17 \pm 0.12$ | 99.932 |
| | 5 min | $3.67 \pm 0.21$ | 99.979 |
| | 15 min | $4.00 \pm 0.10$ | 99.990 |

Shading indicates data points where virucidal efficacy is >99.9% (3 $\text{log}_{10}$ reduction) vs. virus control; virus control=untreated virus, 0 mg/mL astodrimmer sodium SD=standard deviation

**Figure 4.**
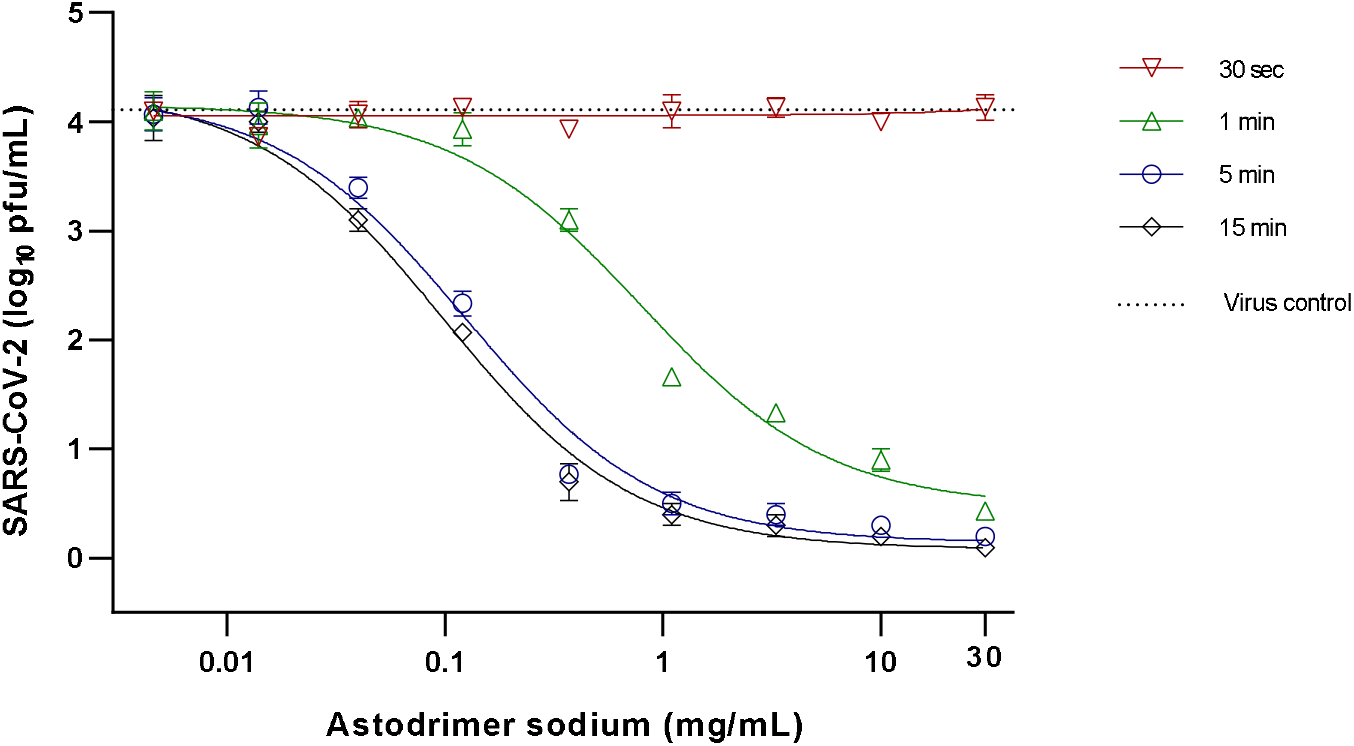
Virucidal efficacy of astodrimer sodium against SARS-CoV-2 (2019-nCoV/USA-WA1/2020) measured by a reduction in mean infectious virus (Log_10_ pfu/mL), at 16 hours post-infection in Vero E6 cells. Astodrimer sodium (0.0046 to 30 mg/mL) was incubated with SARS-CoV-2 (2019-nCoV/USA-WA1/2020) for 30 sec, 1 min, 5 min and 15 min. Treated virus was added to Vero E6 cells and the amount of infectious virus in the supernatant was determined by plaque assay 16 hours post-infection. Graph shows dose-response of astodrimer sodium virucidal activity using 10^4^ pfu/mL virus inoculum. Points and error bars represent mean ± SD of triplicate readings. Dotted line indicates level of mean infectious virus when untreated virus was added to Vero E6 cells (virus control).

### 3.4 Antiviral Efficacy in Primary Human Airway Epithelial Cells

To determine the ability of astodrimer sodium to prevent SARS-CoV-2 infection of primary human epithelial cells, the compound was evaluated against the 2019-nCoV/USA-WA1/2020 strain in HBEpC cell culture. Astodrimer sodium was found to reduce infection of HBEpC primary cells by SARS-CoV-2 by up to 98% vs virus control by nucleocapsid ELISA (Figure 5A), and by up to 95% in the plaque assay (data not shown). In contrast, treatment with iota-carrageenan had minimal antiviral effect against SARS-CoV-2 in this cell line, with the highest concentration tested reducing infection by just 17% by nucleocapsid ELISA (Figure 5B), and just 21% in the plaque assay (data not shown). The maximum level of inhibition with astodrimer sodium was comparable to inhibition achieved with the SARS-CoV-2 spike protein antibody (pAb, T01KHuRb) positive control (see Figure 5A and B).

**Figure 5.**
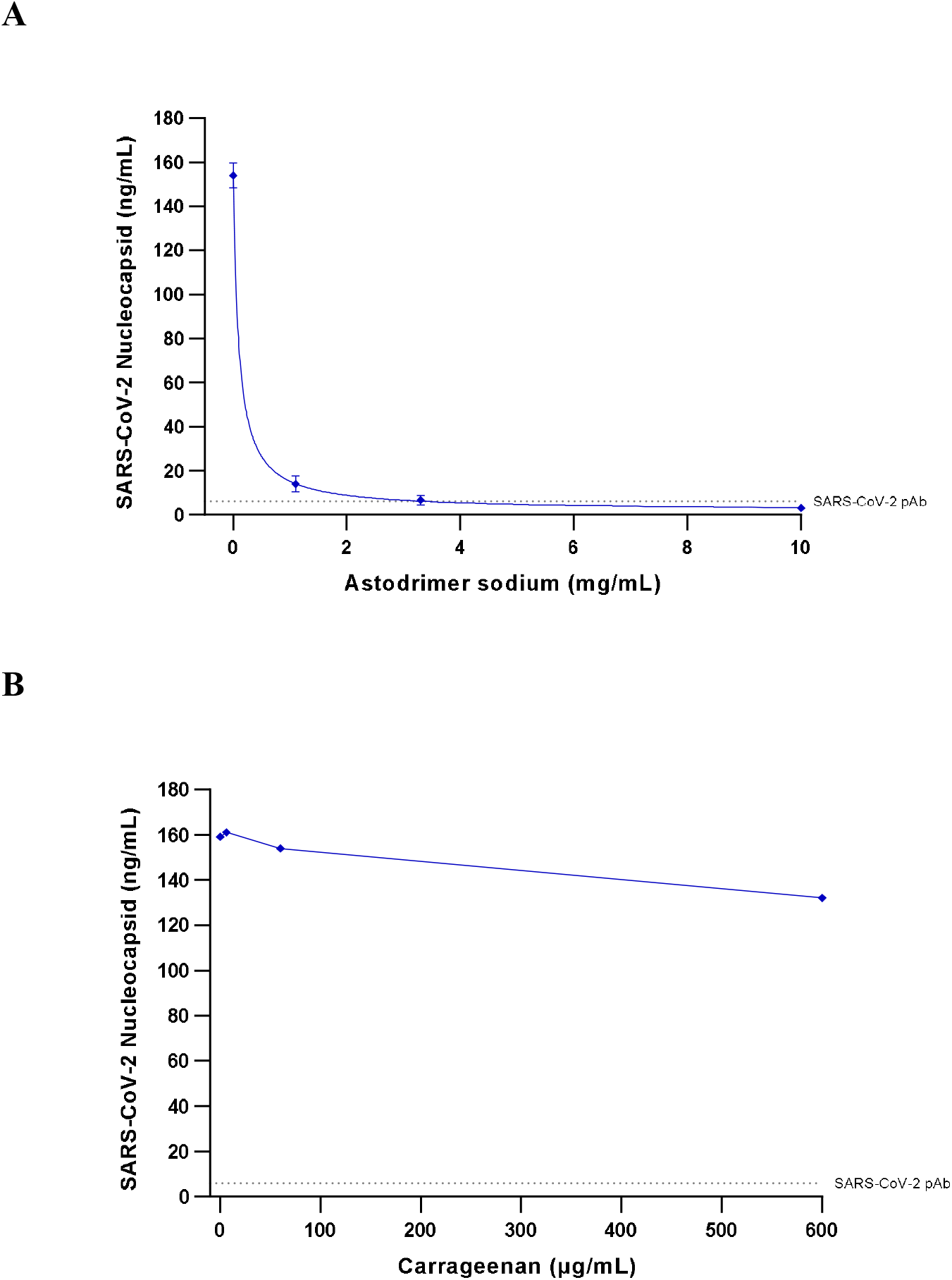
Antiviral efficacy of astodrimer sodium and iota-carrageenan against SARS-CoV-2 (2019-nCoV/USA-WA1/2020) measured by a reduction in nucleocapsid (ng/mL), at Day 4 post-infection in human bronchial epithelial primary cells (HBEpC) Astodrimer sodium (0, 1.1, 3.3 and 10 mg/mL) or iota-carrageenan (0, 6, 60 and 600 μg/mL) were added to cell cultures 1 hour prior to infection. **A.** Dose-response of astodrimer sodium antiviral activity. Points and error bars represent mean ± SD of triplicate readings. **B.** Dose-response of carrageenan antiviral activity. Points represent one replicate. Dotted lines indicates level of inhibition achieved with positive control, SARS-CoV-2 pAb.

## 4. Discussion

Astodrimer sodium demonstrated potent antiviral activity against globally diverse SARS-CoV-2 strains *in vitro*. Antiviral activity was demonstrated by reduction in CPE, release of infectious virus and release of viral nucleocapsid protein. Antiviral activity was demonstrated when astodrimer sodium was added to cells prior to infection of cells and when the compound was added to cells already exposed to SARS-CoV-2. Irreversible virucidal activity was demonstrated when astodrimer sodium was mixed with virus for as little as 1 minute.

Of note is a significantly high SI for astodrimer sodium in the antiviral assays relative to other antiviral compounds under investigation for SARS-CoV-2 activity (Pizzorno et al., 2020).

Remdesivir was used as the antiviral positive control for the CPE inhibition and antiviral assays and the experimental EC_50_ was consistent with published data generated with a different clinical isolate of SARS-CoV-2 (Wang et al., 2020).

Astodrimer sodium inhibited infection of a human airway primary epithelial cell by SARS-CoV-2, whereas iota-carrageenan, which is a polyanionic compound in marketed nasal spray formulations, failed to provide significant inhibition at concentrations that have previously been shown to reduce SARS-CoV-2 infection in Vero E6 cells (Bansal et al., 2020). The unique structure of astodrimer sodium, a sulphonated, roughly spherical molecule with a core and densely packed branches radiating out from the core, appears to provide potential benefits over other polyanionic compounds such as iota-carrageenan and heparin, which are linear sulphated molecules with a distribution of molecular weight. The authors are not aware of data showing that iota-carrageenan is virucidal, while heparin has demonstrated a lack of irreversible, virucidal interaction with HSV virion components (Ghosh et al., 2009).

The antiviral data are consistent with astodrimer sodium being a potent inhibitor of early events in the virus lifecycle. The virucidal assay data suggest that astodrimer sodium antiviral activity was consistent with the proposed mechanism of action of binding to virus, thereby irreversibly inactivating virus and blocking infection.

The virucidal activity of astodrimer sodium demonstrated that it irreversibly inhibits the early phase of virus infection and replication. These findings suggest potent inhibition of viral attachment, fusion and entry of the virus, which prevents virus replication and release of infectious virus progeny.

Astodrimer sodium has been previously found to be an effective antiviral that exerts its inhibition in the early virus-host receptor recognition interactions (Tyssen et al., 2010; Telwatte et al., 2011), and its potential mechanism of action against SARS-CoV-2 is likely similar to that identified for other pathogens. Astodrimer sodium was found to bind to HIV-1 by strong electrostatic forces to positively charged clusters of highly conserved amino acids on HIV-1 gp120 protein and/or positively charged amino acid regions located between the stems of V1/V2 and V3 loops, which are exposed by conformational changes to gp120 after viral binding to the receptor/co-receptor complex (Tyssen et al., 2010; Connell and Lortat-Jacob, 2013).

Many viruses utilize negatively charged heparan sulfate proteoglycans (HS) on the cell plasma membrane as an initial means to scan the surface of the cell, and to attach in order to chaperone the virus onto the receptor complex prior to viral entry (Sarrazin et al., 2011; Connell and Lortat-Jacob, 2013). The receptor interactions occur in a sequential manner with virus-HS interactions preceding receptor/co-receptor binding, which combined leads to fusion of the viral envelope and the cell membrane.

Data indicate that astodrimer sodium-gp120 interaction may physically block initial HIV-1 association with HS and thereby block the subsequent virus-receptor complex functions. Virucidal studies of astodrimer sodium determined that it did not disrupt the HIV-1 particle or cause the loss of gp120 spike protein from the viral surface (Telwatte et al., 2011).

A report by Liu et al., 2020, described that densely glycosylated trimeric SARS-CoV-2 spike (S) protein subunit S1, which is important for receptor binding, binds to HS. To engage with the ACE2 receptor, the S protein undergoes a hinge-like conformational change that transiently hides or exposes the determinants of receptor binding (Wrapp et al., 2020). Recent studies have identified the binding of heparin to the receptor binding domain (RBD) of S1 resulting in a conformational change to the S protein (Mycroft-West et al., 2020a, b and c). Mutations in the S protein that are distal from the RBD also impact on viral transmission (Walls et al., 2020; Korber et al., 2020; Yuan et al., 2020). Non-RBD polybasic cleavage sites, including S1/S2 loop (Hoffmann et al., 2020a), have been described on SARS-CoV-2 S protein (Qiao and Olvera de la Cruz, 2020) and may also be a site of potential interaction with astodrimer sodium.

SARS-CoV-2 utilizes the ACE2 receptor for viral infection of host cells (Hoffmann et al., 2020b). Human CoV-NL63 also utilizes HS and ACE2 as its cellular receptor complex (Milewska et al., 2014). The importance of HS for viral infectivity was also demonstrated for close genetically related pseudo-typed SARS-CoV (Lang et al., 2011). The potential dependence of SARS-CoV-2 on HS for attachment and entry combined with antiviral data from other viruses suggest that negatively charged astodrimer sodium may have antiviral activity against SARS-CoV-2 *in vitro* by blocking the early virus-receptor recognition events.

Astodrimer sodium is a polyanionic dendrimer reformulated for use as a topical, nasally administered antiviral agent to inactivate SARS-CoV-2 before infection can occur. The potential advantages of astodrimer sodium over other technologies include its lack of systemic absorption following topical application (Chen et al., 2009; O’Loughlin et al., 2010; McGowan et al., 2011). In addition, the SI of astodrimer sodium for SARS-CoV-2 is high and in a vaginal gel formulation (10 mg/mL), the compound has been shown to be safe and effective in phase 2 and large phase 3 trials for treatment and prevention of BV (Chavoustie et al., 2020; Waldbaum et al., 2020; Schwebke et al., 2021) and is now marketed in Europe, Australia, New Zealand and several countries in Asia. Astodrimer sodium is also the active antiviral substance in VivaGel^®^ condom products that have marketing authorization in Europe, Japan, Australia/New Zealand and Canada. However, these current formulations are not appropriate for use to protect the respiratory tract from SARS-CoV-2 infection.

## 5. Conclusions

Data from the current studies, taken together with studies of astodrimer sodium antiviral activity against HIV-1, and HSV-1 and −2, indicate that the compound exerts its antiviral activity against geographically diverse SARS-CoV-2 isolates by interfering with the early virus-cell recognition events. Astodrimer sodium is a potent virucidal agent that reduces the infectivity of SARS-CoV-2 by −99.9% after 1 minute of exposure to the virus. These studies support astodrimer sodium being able to prevent early virus entry steps such as attachment, thereby reducing or preventing viral infection or cell-cell spread.

An antiviral agent such as astodrimer sodium that blocks binding of the virus to target cells could potentially be used as a preventive and/or a therapeutic agent against SARS-CoV-2. These antiviral studies suggest that reformulation of astodrimer sodium for delivery to the respiratory tract may be an effective preventive strategy to block SARS-CoV-2 transmission and augment other protective and therapeutic strategies.

The potent antiviral and virucidal activity of astodrimer sodium against SARS-CoV-2 warrants further investigation.

---

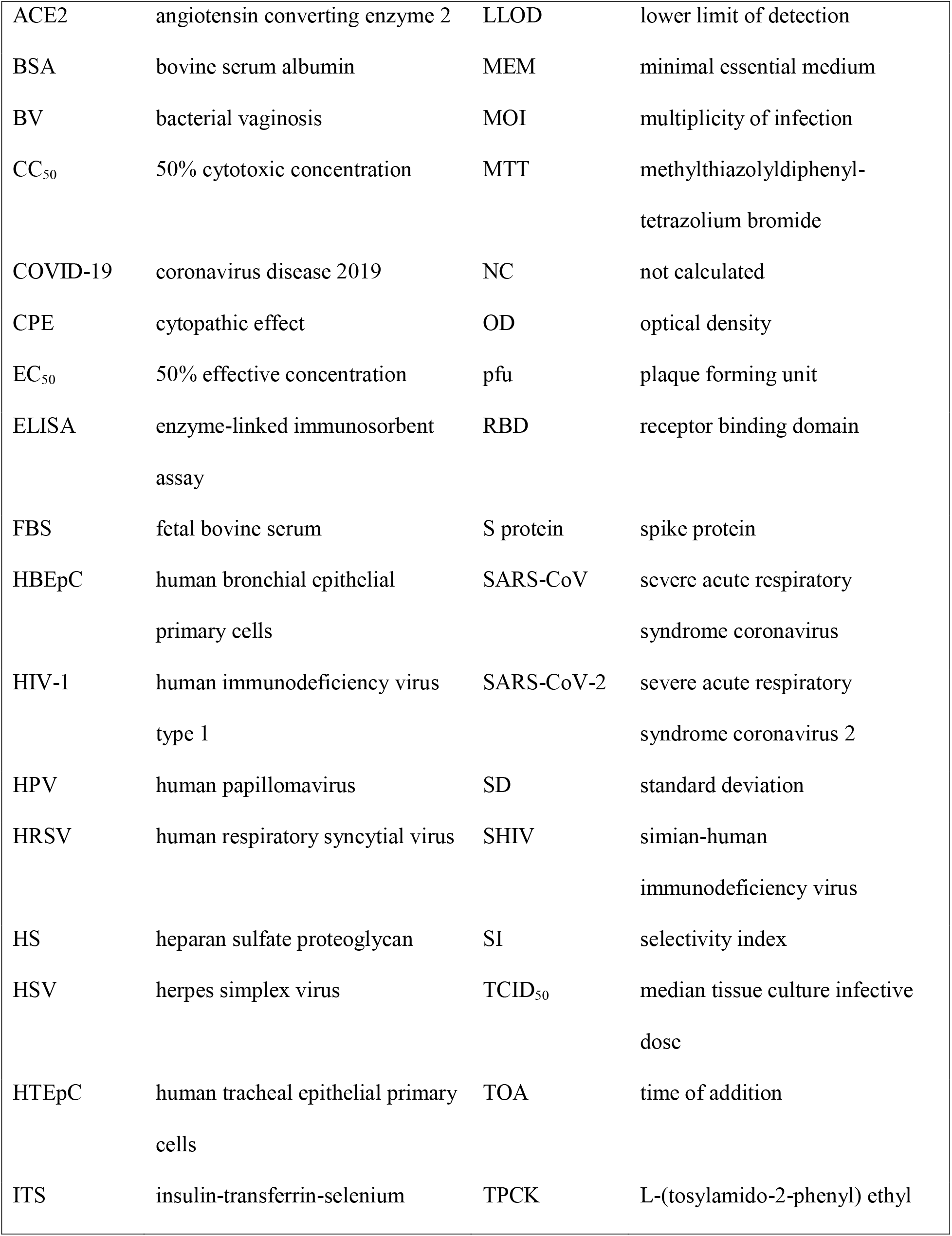

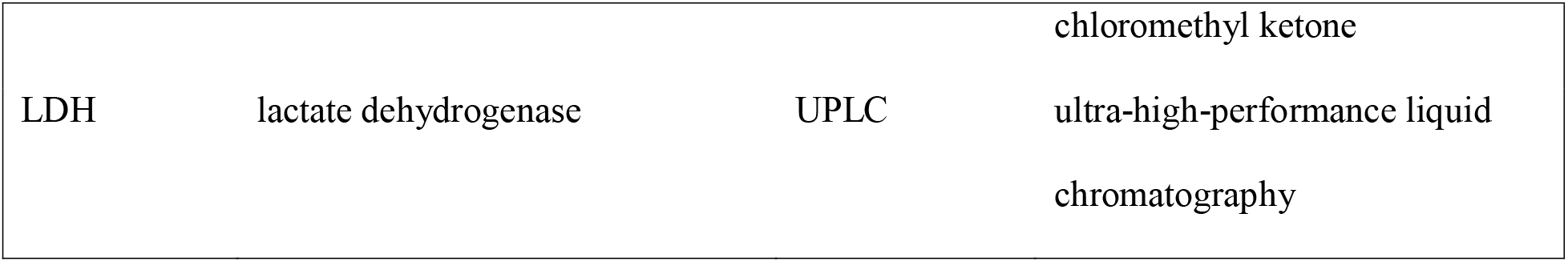

## Acknowledgments

The authors were fully responsible for the content, editorial decisions, and opinions expressed in the current article. The authors would like to acknowledge 360Biolabs Pty Ltd (Melbourne, Australia) for the conduct of the CPE assays and Scripps Research Institute for the conduct of the antiviral and virucidal assays. The authors would also like to thank the Peter Doherty Institute for Infection and Immunity (Melbourne, Australia) for the gift to 360Biolabs of SARS-CoV-2 hCoV-19/Australia/VIC01/2020 used in these studies.

